# A comprehensive analysis of 3’ end sequencing data sets reveals novel polyadenylation signals and the repressive role of heterogenous ribonucleoprotein C on cleavage and polyadenylation

**DOI:** 10.1101/033001

**Authors:** Andreas J. Gruber, Ralf Schmidt, Andreas R. Gruber, Georges Martin, Souvik Ghosh, Manuel Belmadani, Walter Keller, Mihaela Zavolan

## Abstract

Alternative polyadenylation (APA) is a general mechanism of transcript diversification in mammals, which has been recently linked to proliferative states and cancer. Different 3’ untranslated region (3’ UTR) isoforms interact with different RNA binding proteins (RBPs), which modify the stability, translation, and subcellular localization of the corresponding transcripts. Although the heterogeneity of pre-mRNA 3’ end processing has been established with high-throughput approaches, the mechanisms that underlie systematic changes in 3’ UTR lengths remain to be characterized. Through a uniform analysis of a large number of 3’ end sequencing data sets we have uncovered 18 signals, 6 of which novel, whose positioning with respect to pre-mRNA cleavage sites indicates a role in pre-mRNA 3’ end processing in both mouse and human. With 3’ end sequencing we have demonstrated that the heterogeneous ribonucleoprotein C (HNRNPC), which binds the poly(U) motif whose frequency also peaks in the vicinity of polyadenylation (poly(A)) sites, has a genome-wide effect on poly(A) site usage. HNRNPC-regulated 3’ UTRs are enriched in ELAV-like RNA binding protein 1 (ELAVL1) binding sites and include those of the CD47 molecule (CD47) gene, which participate in the recently discovered mechanism of 3’ UTR-dependent protein localization (UDPL). Our study thus establishes an up-to-date, high-confidence catalog of 3’ end processing sites and poly(A) signals and it uncovers an important role of HNRNPC in regulating 3’ end processing. It further suggests that U-rich elements mediate interactions with multiple RBPs that regulate different stages in a transcript’s life cycle.

## Introduction

The 3’ ends of most RNA polymerase ll-generated transcripts are generated through endonucleolytic cleavage and addition of a poly(A) tail of 70–100 nucleotides median length (Subtelny et al. 2014). Recent studies have revealed systematic changes in 3’ UTR lengths upon changes in cellular states, either physiological (Sandberg et al. 2008; Berg et al. 2012) or during pathologies (Masamha et al. 2014). 3’ UTR lengths are sensitive to the abundance of specific spliceosomal proteins (Kaida et al. 2010), core pre-mRNA 3’ end processing factors (Martin et al. 2012; Gruber et al. 2012), and polyadenylation factors (Jenal et al. 2012). Because 3’ UTRs contain many recognition elements for RNA-binding proteins that regulate the subcellular localization, intra-cellular traffic, decay and translation rate of the transcripts in different cellular contexts (see e.g. Nam et al. 2014), the choice of polyadenylation (poly(A)) sites has important regulatory consequences that reach up to the sub-cellular localization of the resulting protein (Berkovits and Mayr 2015). Studies of presumed regulators of polyadenylation would greatly benefit from the general availability of comprehensive catalogs of poly(A) sites such as PolyA_DB (Zhang et al. 2005; Lee et al. 2007), which was introduced in 2005 and updated two years later.

Full-length cDNA sequencing offered a first glimpse on the pervasiveness of transcription across the genome and on the complexity of gene structures (Kawai et al. 2001). Next-generation sequencing technologies, frequently coupled with the capture of transcript 5’ or 3’ ends with specific protocols, enabled the quantification of gene expression and transcript isoform abundance (Katz et al. 2010). By increasing the depth of coverage of transcription start sites and mRNA 3’ ends, these protocols aimed to improve the quantification accuracy (de Hoon and Hayashizaki 2008; Ozsolak et al. 2009; Beck et al. 2010; Shepard et al. 2011). Sequencing of mRNA 3’ ends takes advantage of the poly(A) tail, which can be captured with an oligo-dT primer. Over 4.5 billion reads were obtained with several protocols from human or mouse mRNA 3’ ends in a variety of cell lines (Shepard et al. 2011; Lin et al. 2012), tissues (Derti et al. 2012; You et al. 2014), developmental stages (Ulitsky et al. 2012; Li et al. 2012), cell differentiation stages (Hoque et al. 2013) as well as following perturbations of specific RNA processing factors (Martin et al. 2012; Gruber et al. 2012; Jenal et al. 2012; Almada et al. 2013; Ji et al. 2013). Although some steps are shared by many of the proposed 3’ end sequencing protocols, the studies that employed these methods have reported widely varying numbers of 3’ end processing sites. For example, between 54,686 (Lee et al. 2007), 439390 (Derti et al. 2012) and 1,287,130 (Lin et al. 2012) sites have been reported in the human genome.

The current knowledge about sequence motifs that are relevant for pre-mRNA cleavage and polyadenylation, (for review see (Proudfoot 2011)), goes back to studies conducted before next generation sequencing technologies became broadly used (Proudfoot and Brownlee 1976; Beaudoing et al. 2000; Tian et al. 2005). These studies revealed that the AAUAAA hexamer, which recently was found to bind the WDR33 and CPSF4 subunits of the cleavage and polyadenylation specificity factor (CPSF) (Schönemann et al. 2014; Chan et al. 2014) as well as some close variants, are highly enriched upstream of the pre-mRNA cleavage site. The A[AU]UAAA *cis*-regulatory element (also called poly(A) signal) plays an important role in pre-mRNA cleavage and polyadenylation (Tian and Graber 2012) and is located upstream of a large proportion of pre-mRNA cleavage sites identified in different studies (Graber et al. 1999; MacDonald and Redondo 2002; Tian et al. 2005). However, some transcripts that do not have this poly(A) signal are nevertheless processed, indicating that the poly(A) signal is not absolutely necessary for cleavage and polyadenylation. The constraints that functional poly(A) signals have to fulfill are not entirely clear, and at least ten other hexamers have been proposed to have this function (Beaudoing et al. 2000).

Viral RNAs as for example from the simian virus 40 have been instrumental in uncovering RNA binding protein (RBP) regulators of polyadenylation and their corresponding sequence elements. Early studies revealed modulation of poly(A) site usage by U-rich element binding proteins such as the heterogeneous nuclear ribonucleoprotein (hnRNP) C1/C2 (Wilusz et al. 1988; Zhao et al. 2005), the polypyrimidine tract binding protein 1 (Castelo-Branco et al. 2004; Zhao et al. 2005), Fip1L1 and CSTF2 (Zhao et al. 2005), while more recent studies described the modulation of poly(A) site usage by proteins that bind G-rich elements - cleavage stimulation factor CSTF2 (Alkan et al. 2006) and HNRNPs F and H1 (Arhin et al. 2002) - or C-rich elements - poly(rC)-binding protein 2(Ji et al. 2013). Some of these proteins are multi-functional splicing factors that appear to couple various steps in pre-mRNA processing, such as splicing, cleavage and polyadenylation (Millevoi et al. 2009). The sequence elements to which these regulators bind are also frequently multi-functional, enabling positive or negative regulation by different RBPs (Alkan et al. 2006). A first step towards understanding the regulation of poly(A) site choice is to construct genome-wide maps of poly(A) sites, which can be used to investigate differential polyadenylation across tissues and the response of poly(A) sites to specific perturbations.

To this end, we integrated the majority of the large-scale 3’ end sequencing data sets that are available and generated “PolyAsite” (http://www.polyasite.unibas.ch). To further illustrate the power of this resource, we updated the set of poly(A) signals, identifying sequence motifs whose positioning with respect to the pre-mRNA cleavage sites suggests an involvement in polyadenylation. Our analysis not only recovered the twelve motifs that were identified previously (Beaudoing et al. 2000) but revealed six additional signals that have a similar characteristic positional preference in both human and mouse. Among the motifs that are enriched around poly(A) sites but have a positional preference different from that of poly(A) signals we found the poly(U) motif, which is bound by a variety of proteins, including the splicing-related factor HNRNPC. With CLIP we confirmed that HNRNPC indeed binds in the vicinity of poly(A) sites and with siRNA-mediated HNRNPC knock-down we demonstrated that HNRNPC target regions behave in a manner indicative of HNRNPC masking poly(A) sites and protecting nascent transcripts from cleavage and polyadenylation. Finally, we found that the relative abundance of CD47 3’ UTR isoforms, whose interaction with the U-rich element binding protein ELAVL1 results in differential localization of the encoded protein between endoplasmic reticulum and plasma membrane (Berkovits and Mayr2015), is regulated by HNRNPC.

## Results

### Preliminary processing of 3’ end sequencing data sets

Protocol specific biases as well as vastly different computational data processing strategies may explain the discrepancy in the reported number of 3’ end processing sites, which ranges from less than one hundred thousand to over one million (Derti et al. 2012; Lin et al. 2012; Lee et al. 2007) for the human genome. Comparing the 3’ end processing sites from two recent genome-wide studies (Derti et al. 2012; You et al. 2014) we found that a substantial proportion was unique to one or the other of the two studies (Supplementary Table 1). This motivated us to develop a uniform and flexible processing pipeline that facilitates the incorporation of all published sequencing data sets yielding a comprehensive set of high-confidence 3’ end processing sites. From public databases we obtained 78 human and 110 mouse data sets of 3’ end sequencing reads (Supplementary Tables 2 and 3), generated with 9 different protocols, for which sufficient information to permit the appropriate pre-processing steps (trimming of 5’ and 3’ adaptor sequences, reverse-complementing the reads, etc., as appropriate) was available. We pre-processed each sample as appropriate given the underlying protocol and then subjected all data sets to a uniform analysis as follows. We mapped the pre-processed reads to the corresponding genome and transcriptome and identified unique putative 3’ end processing sites. Because many protocols employ oligo-dT priming to capture the pre-mRNA 3’ ends, internal priming is a common source of false positive sites, which we tried to identify and filter out as described in the Methods section. From the nearly 200 3’ end sequencing libraries we thus obtained an initial set of 6,983,499 putative 3’ end processing sites for human and 8,376,450 for mouse. The majority of these sites (76% for human and 71% for mouse) has support in only one sample, consistent with our initial observations of limited overlap between the sets of sites identified in individual studies, and mirroring also the results of transcription start site mapping with the CAGE technology (FANTOM Consortium and the RIKEN PMI and CLST (DGT) et al. 2014). Nevertheless, we developed an analysis protocol that aimed to identify *bona fide*, independently regulated poly(A) sites, including those that have been captured in a single sample. To do this, we used not only the sequencing data, but also information about poly(A) signals, which we therefore set to comprehensively identify in the first step of our analysis.

### Highly specific positioning with respect to the pre-mRNA cleavage site reveals novel polyadenylation signals

To search for signals that may guide polyadenylation, we designed a very stringent procedure to identify high-confidence 3’ end processing sites. Pre-mRNA cleavage is not completely deterministic, but occurs with higher frequency at “strong” 3’ end processing sites and with low frequency at neighboring positions (Tian et al. 2005). Therefore, a common step in the analysis of 3’ end sequencing data is to cluster putative sites that are closely spaced and report the dominant site from each cluster (Tian et al. 2005; Martin et al. 2012; Lianoglou et al. 2013). To determine an appropriate distance threshold, we ranked all the putative sites first by the number of samples in which they were captured and then by the normalized number of reads in these samples. Traversing the list of sites from those with the strongest to those with weakest support, we associated lower-ranking sites located up to a specific distance from the higher-ranked site with the corresponding higher-ranking site. We scanned the range of distances of [0, 25] nucleotides (nts) upstream and downstream of the high-ranking site and we found that the proportion of putative 3’ end processing sites that are merged into clusters containing more than one site reaches 40% at ~8 nts, and changes little by further increasing the distance (see also “Clustering of closely spaced 3' end sites into 3' end processing regions” in the Methods section). For consistency with previous studies (Tian et al. 2005), we used a distance of 12 nts. To reduce the frequency of protocol-specific artifacts, we used only clusters that were supported by reads derived with at least 2 protocols, and to allow unambiguous association of signals to clusters we only used for the signal inference clusters that did not have another cluster within 60 nucleotides. This procedure resulted in 221,587 3’ end processing clusters for human and 209,345 for mouse.

Analyzing 55 nt-long regions located immediately upstream of the center of these 3’ end processing clusters (as described in the Methods section) we found that the canonical poly(A) signals AAUAAA and AUUAAA are highly enriched and have a strong positional preference, peaking at 21 nts upstream of cleavage sites (Figure 1A), as reported previously (Beaudoing et al. 2000; Tian et al. 2005). We therefore asked whether other hexamers have a similarly peaked frequency profile, which would be indicative of their functioning as poly(A) signals. The twelve signals that were identified in a previous study (Beaudoing et al. 2000) served as controls for the procedure. In both mouse and human data, the motif with the highest peak was, as expected, the canonical poly(A) signal, AAUAAA, which occurred in 46.82% and 39.54% of the human and mouse sequences, respectively. Beyond this canonical signal, we found 21 additional hexamers, the second most frequent being the close variant of the canonical signal, AUUAAA, which was present in 14,52% and 12,28% of the human and mouse 3’ sequences, respectively. All twelve known polyadenylation signals (Beaudoing et al. 2000) were recovered by our analysis in both species, demonstrating the reliability of our approach. Further supporting this conclusion is the fact that six of the ten newly identified signals that we identified in each of the two species are also shared. All of the conserved signals are very close variants (one nt difference except for AACAAG) of one of the two main poly(A) signals, AAUAAA and AUUAAA. Strikingly, all of these signals peak in frequency at 20–22 upstream of the cleavage site (Figure 1A). Experimental evidence for single nucleotide variants of the AAUAAA signal (including the AACAAA, AAUAAU, and AAUAAG motifs identified here) functioning in polyadenylation was already provided by Sheets et al. (Sheets et al. 1990). The four signals identified in only one of each species also had a clear peak at the expected position with respect to the poly(A) site, but they had a larger variance (Supplementary Figure 1). Altogether, these results indicate a genuine role of the newly identified signals in the process of cleavage and polyadenylation.

**Figure 1.**
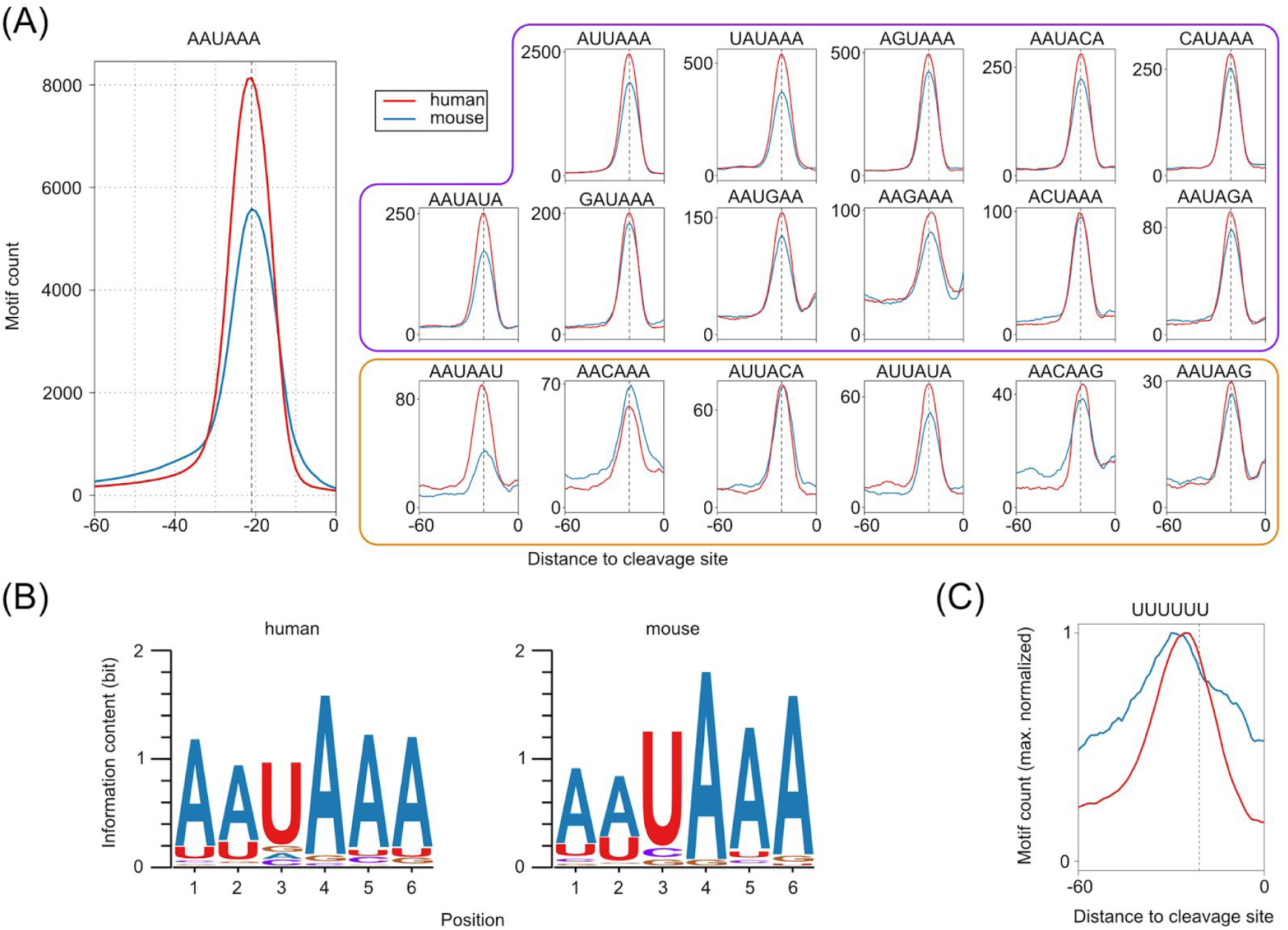
Hexamers with highly specific positioning upstream of human and mouse pre-mRNA 3’ end cleavage sites –. **(A)** The frequency profiles of the 18 hexamers that show a specific positional preference, as expected for poly(A) signals, in both human and mouse. The known poly(A) signal, AAUAAA, has the highest frequency of occurrence (left panel). Apart from the twelve signals previously identified (Beaudoing et al. 2000), AAUAAA and motifs with the purple frame, we have identified six additional motifs (orange frame) whose positional preference with respect to poly(A) sites suggests that they function as poly(A) signals and are conserved between human and mouse. **(B)** Sequence logos based on all occurrences of the eighteen motifs upstream of the poly(A) sites from the human (left) and mouse (right) atlas. **(C)** The (U)_6_ motif, which is also enriched upstream of pre-mRNA cleavage sites, has a broader frequency profile and peaks upstream of the poly(A) signals, which are precisely positioned 20–22 nucleotides upstream of the pre-mRNA cleavage sites (indicated by the dashed, vertical line).

87% and 79%, respectively, of the 221,587 high-confidence 3’ end processing clusters in human and 209,345 in mouse, had at least one of the 22 signals identified above in their upstream region. Even when considering only the eighteen signals that are conserved between human and mouse, 86% of the human clusters and 75% of the mouse clusters had a poly(A) signal. Thus, our analysis almost doubles the set of poly(A) signals and suggests that the vast majority of poly(A) sites does indeed have a poly(A) signal which is positioned very precisely with respect to the pre-mRNA cleavage site. The dominance of the canonical poly(A) signal is reflected in the sequence logos constructed based on all annotated hexamers in the human and mouse poly(A) site atlases, as will be described in the following section and in the Methods section (see “Sequence logos of the identified poly(A) signals”) (Figure 1B).

### A comprehensive catalog of high-confidence 3’ end processing sites

Based on all of the 3’ end sequencing data sets available and the conserved poly(A) signals that we inferred as described above, we constructed a comprehensive catalog of strongly supported 3’ end processing sites in both mouse and human genomes (see “Supplementary Materials” for more details about the protocols that were used to generate these data sets). We started from the 6,983,499 putative cleavage sites for human and 8,376,450 for mouse derived as described in section “Preliminary processing of 3’ end sequencing data sets”. Although in many data sets a large proportion of putative sites was supported by single reads and did not have any of the expected poly(A) signals in the upstream region, the incidence of upstream poly(A) signals increased with the number of reads supporting a putative site (Supplementary Figure 2). Thus, we used the frequency of occurrence of poly(A) signals to define sample-specific cutoffs for the number of reads required to support a putative cleavage site, as described in the Methods section. We then clustered all putative sites with sufficient read support, associating lower ranked sites with higher-ranking sites that were located within at most 12 nucleotides upstream or downstream, as described above (see “Highly specific positioning with respect to the pre-mRNA cleavage site reveals novel polyadenylation signals”). Because in this set of clusters we found cases where the pre-mRNA cleavage site appeared located in an A-rich region upstream of another putative cleavage site, we specifically reviewed clusters in which a putative cleavage site was very close to a poly(A) signal, as these likely reflect internal priming events (Gruber et al. 2014; Derti et al. 2012; Shepard et al. 2011). These clusters were either associated with a downstream cluster, retained as independent clusters, or discarded, according to the procedure outlined in the Methods section (see “Treatment of putative 3’ end sites originating from internal priming”). Reasoning that distinct 3’ end processing sites should have independent signals to guide their processing, we merged clusters that shared all poly(A) signal(s) within 60 nucleotides upstream of their representative sites, clusters whose combined span was less than 25 nucleotides, and clusters without annotated poly(A) signals that were closer than 12 nucleotides to each other and had a combined span of at most 50 nucleotides. Clusters without poly(A) signals and larger than 50 nucleotides were excluded from the atlas. This procedure (for details see the Methods section) resulted in 392,912 human and 183,225 mouse 3’ end processing clusters. Of note, even though 3’ end processing sites that were within 25 nucleotides of each other were merged into single clusters, the median cluster span was very small, 7 and 3 nucleotides for mouse and human, respectively (Supplementary Figure 3). Supplementary Figures 4A and 5A show the frequency of occurrence of the four nucleotides as a function of the distance to the cleavage sites, for sites that were supported by a decreasing number of protocols. These profiles exhibit the expected pattern (Tian et al. 2005; Ozsolak et al. 2010; Martin et al. 2012), indicating that our approach identified *bona fide* 3’ end processing sites, even when they had limited experimental support.

The proportion of clusters located in the terminal exon increased with an increasing level of support (Supplementary Figures 4B and 5B), probably indicating that the canonical poly(A) sites of constitutively expressed transcripts are identified by the majority of protocols, whereas poly(A) sites that are only used in some conditions were captured only in a subset of experiments. Although in constructing our catalog we used most of the reads generated in two recent studies (You et al. 2014; Derti et al. 2012) (more than 95% of the reads that supported human 3’ end processing sites in these two data sets mapped within the poly(A) site clusters of our human catalog), only 61.82% (You et al. 2014) and 41.38% (Derti et al. 2012) of the unique processing sites inferred in these studies were located within poly(A) clusters from our human catalog. This indicates that a large fraction of the sites that were catalogued in previous studies is supported by a very small number of reads and lack canonically positioned poly(A) signals. We applied very stringent rules to construct an atlas of high-confidence poly(A) sites and the entire set of putative cleavage sites that resulted from mapping all of the reads obtained in these 3’ end sequencing studies is available online at http://www.polyasite.unibas.ch, where users can filter sites of interest based on the number of supporting protocols, the identified poly(A) signals, or the genomic context of the clusters.

### 3’ end processing regions are enriched in poly(U)

76% and 75% of the human and mouse 3’ end processing sites from our poly(A) atlases, respectively, possessed a conserved poly(A) signal in their 60 nt upstream region. That ~25% did not may reflect the possibility that pre-mRNA cleavage and polyadenylation does not absolutely require a poly(A) signal (Venkataraman et al. 2005). Nevertheless, we asked whether these sites possess other signals, with a different positional preference, which may contribute to their processing. To answer this question, we searched for hexamers that were significantly enriched in the 60 nt upstream of clusters without an annotated poly(A) signal. The two most enriched hexamers were poly(A) (p-value of binomial test < 1.0e-100 (see “Hexamer enrichment in upstream regions of 3’ end clusters”)), which showed a broad peak in the region of -20 to -10 upstream of cleavage sites, and poly(U) (p-value < 1.0e-100), which also has a broad peak around -25 nts upstream of cleavage sites, particularly pronounced in the human data set (Figure 1C). The poly(U) hexamer is very significantly enriched (p-value of binomial test < 1.0e-100) in the 60 nts upstream regions of all poly(A) sites, not only in those that do not have a common poly(A) signal (11th most enriched hexamer in the human atlas and 60th most enriched hexamer in the mouse atlas, Supplementary Tables 4 and 5, see also “Hexamer enrichment in upstream regions of 3’ end clusters”). Although the A- and U-richness of pre-mRNA 3’ end processing regions have been observed before (Tian and Graber 2012), their relevance for polyadenylation and the regulators that bind these motifs have been characterized only partially. For example, the core 3’ end processing factor FIP1L1 can bind poly(U) (Kaufmann et al. 2004; Lackford et al. 2014) and its knock-down causes a systematic increase in 3’UTR lengths (Lackford et al. 2014; Li et al. 2015).

### HNRNPC knock-down causes global changes in alternative cleavage and polyadenylation

Several proteins (ELAVL1, TIA1, TIAL1, U2AF2, CPEB2 and CPEB4, HNRNPC) that regulate pre-mRNA splicing and polyadenylation, as well as mRNA stability and metabolism have also been reported to bind U-rich elements (Ray et al. 2013). Of these, HNRNPC has been recently studied with individual nucleotide resolution CLIP (iCLIP) and found to bind the majority of protein-coding genes (König et al. 2010), with high specificity for poly(U) tracts (Ray et al. 2013; Liu et al. 2015; König et al. 2010; Cieniková et al. 2014; Zarnack et al. 2013). HNRNPC appears to nucleate the formation of ribonucleoprotein particles on nascent transcripts and to regulate pre-mRNA splicing (Zarnack et al. 2013; König et al. 2010) and polyadenylation at Alu repeats (Tajnik et al. 2015). We therefore hypothesized that HNRNPC binds to the U-rich regions in the vicinity of poly(A) sites and globally regulates not only splicing, but also pre-mRNA cleavage and polyadenylation.

To test this hypothesis, we generated two sets of pre-mRNA 3’ end sequencing libraries from HEK 293 cells that were either transfected with a control siRNA, or an siRNA directed against HNRNPC. The siRNA was very efficient, strongly reducing the HNRNPC protein expression, as shown in Supplementary Figure 6. To evaluate the effect of HNRNPC knock-down on polyadenylation we focused on exons with multiple poly(A) sites. We identified 12,136 such sites with a total of 22,698,094 mapped reads (Supplementary Table 6). We calculated the relative usage of a poly(A) site in a given sample as the proportion of reads that mapped to that site among the reads mapping to any 3’ end processing site in the corresponding exon. We then computed the change in relative usage of each poly(A) site in si-HNRNPC-treated cells compared to control siRNA-treated cells. We found that HNRNPC knock-down affects a large proportion of transcripts with multiple poly(A) sites, reminiscent of what we previously reported for the 25 and 68 kDa subunits of the cleavage factor I (CF I_m_) (Martin et al. 2012; Gruber et al. 2012). To find out whether HNRNPC systematically increases or decreases 3’ UTR lengths, we examined the relative position of poly(A) sites whose usage increases or decreases most strongly in response to HNRNPC knock-down, within 3’ UTRs. The results indicate that poly(A) sites whose usage increases and decreases upon HNRNPC knock-down tend to be located distally and proximally, respectively, within exons (Figure 2A). We confirmed the systematic increase in 3’ UTR lengths upon HNRNPC knock-down by comparing the proximal-to-distal poly(A) site usage ratios of exons that had exactly two polyadenylation sites (Supplementary Figure 7, replicate 1 p-value: 1.1 e-19, replicate 2 p-value: 3.1e-61, one-sided Wilcoxon Rank Sum test). It was noted before that distal poly(A) sites are predominantly used in HEK 293 cells (Martin et al. 2012). Indeed, the proportion of dominant (>50% relative usage) distal sites was 61.75% and 62.58%, respectively, in the two control siRNA-treated samples. However, this proportion increased further in the si-HNRN PC-treated samples, to 64.16% and 65.67%, respectively, consistent with HNRNPC predominantly decreasing the lengths of 3’ UTRs. We therefore concluded that in contrast to CF I_m_, the HNRNPC knock-down causes an overall proximal-to-distal shift in poly(A) site usage.

**Figure 2.**
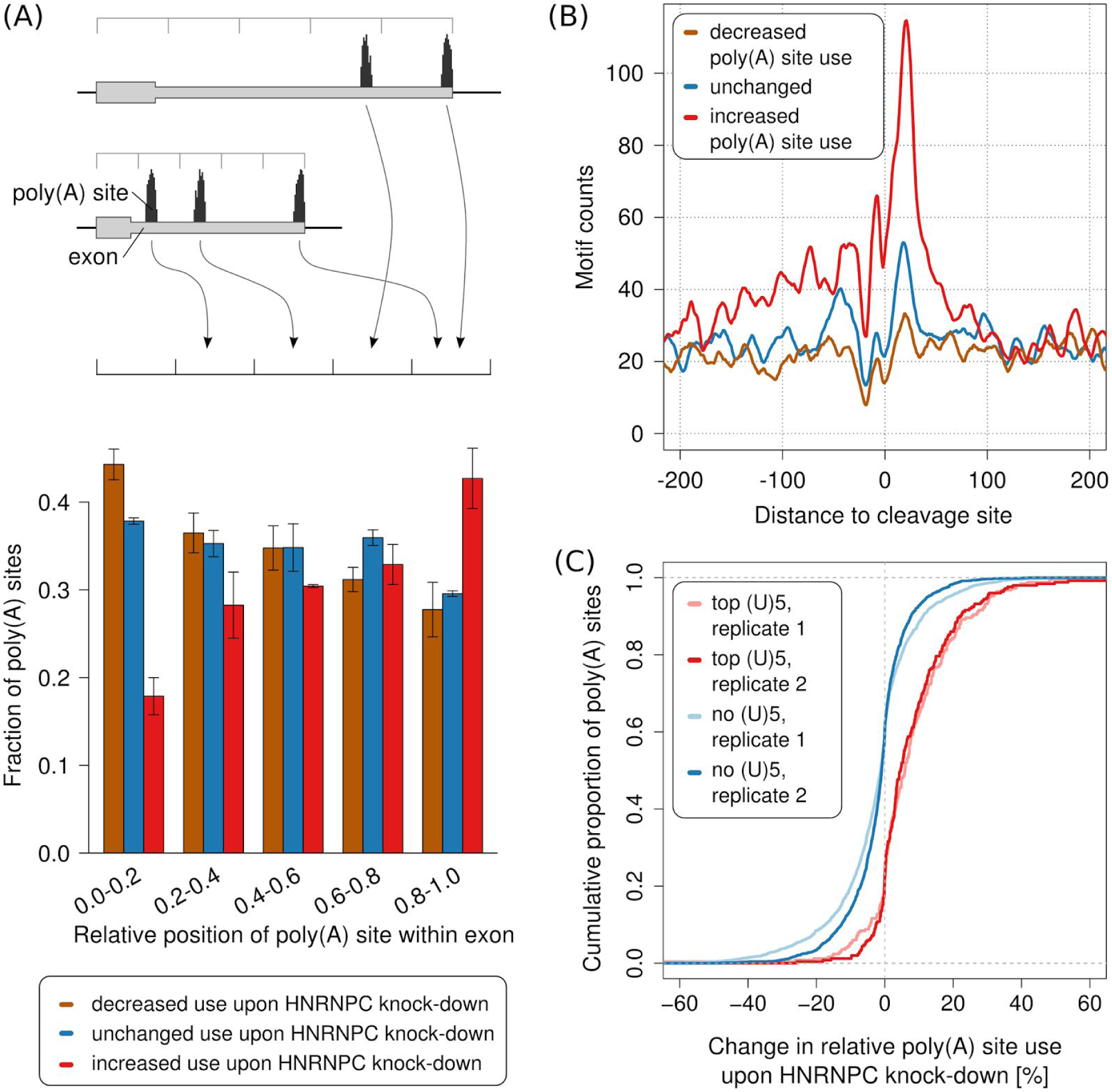
siRNA-mediated knock-down of HNRNPC leads to increased use of distal poly(A) sites. **(A)** Relative location of sites whose usage decreases (brown), does not change (blue) or increases (red) in response to HNRNPC knock-down within 3’ UTRs. We identified the 1000 poly(A) sites whose usage increased most, the 1000 whose usage decreased most and the 1000 whose usage changed least upon HNRNPC knock-down, divided the associated terminal exons into 5 bins, each covering 20% of the exon’s length and computed the fraction of poly(A) sites that corresponded to each of the three categories within each position bin independently. Values represent means and standard deviations from the two replicate HNRNPC knock-down experiments. **(B)** Smoothened (+/-5nt) density of non-overlapping (U)_5_ tracts in the vicinity of sites with a consistent behavior (increased, unchanged, decreased use) in the two HNRNPC knock-down experiments (see “Sequencing of A-Seq2 libraries and quantification of relative poly(A) site usage”). **(C)** Cumulative density function of the percentual change in usage of the 250 poly(A) sites with the highest number of (U)_5_ motifs within +/- 50 nucleotides around their cleavage site (red) and of poly(A) sites that do not contain any (U)_5_ tract within +/- 200 nucleotides (blue), upon HNRNPC knock-down.

As HNRNPC binds RNAs in a sequence-specific manner, one expects an enrichment of HNRNPC binding sites in the vicinity of poly(A) sites whose usage is affected by the HNRNPC knock-down. Indeed, this is what we observed. The density of (U)_5_ tracts, previously reported to be the binding sites of HNRNPC (König et al. 2010; Ray et al. 2013; Liu et al. 2015), is markedly higher around poly(A) sites whose usage increased upon HNRNPC knock-down compared to sites whose relative usage does not change or decreases upon HNRNPC knock-down (Figure 2B). To exclude the possibility that this profile is due to a small number of regions that are very U-rich, we also determined the fraction of poly(A) sites that contained (U)_5_ tracts among the poly(A) sites whose usage increased, decreased or did not change upon HNRNPC knock-down (Supplementary Figure 8). Consistent with the results shown in Figure 2B, we found a higher proportion of (U)_5_ tract-containing poly(A) sites among those whose usage increased upon HNRNPC knock-down compared to those whose usage decreased or was not changed. To further validate HNRNPC binding at the de-repressed poly(A) sites we carried out HNRNPC CLIP and found, indeed, that de-repressed sites have a higher density of HNRNPC CLIP reads compared to other poly(A) sites (Supplementary Figure 9). Finally, we found that poly(A) sites with the highest density of (U)_5_ tracts in the 100 nt region centered on the cleavage site were reproducibly used with increased frequency upon HNRNPC knock-down relative to poly(A) sites that did not contain any binding sites within 200 nucleotides upstream or downstream (Figure 2C; replicate 1 p-value: 2.4e-36, replicate 2 p-value: 1.9e-42, one-sided Mann-Whitney U test). We therefore concluded that HNRNPC’s binding in close proximity of 3’ end processing sites likely masks them from cleavage and polyadenylation.

### Both the number and the length of the uridine tracts contribute to the HNRNPC-dependent poly(A) site usage

If the above conclusions were correct, the effect of HNRNPC knock-down should decrease with the distance between the poly(A) site and the HNRNPC binding sites. Thus we determined the mean change in usage of sites with high densities of poly(U) tracts at different distances with respect to the cleavage site, upon HNRNPC knock-down. As shown in Figure 3A, we found that the largest change in poly(A) site use is observed for poly(A) sites that have a high density of poly(U) tracts in the 100 nt window centered on the cleavage site. The apparent efficacy of HNRNPC binding sites in modulating polyadenylation decreases with their distance to poly(A) sites, and persists over larger distances upstream of the poly(A) site compared to regions downstream of the poly(A) site (Figure 3A).

**Figure 3.**
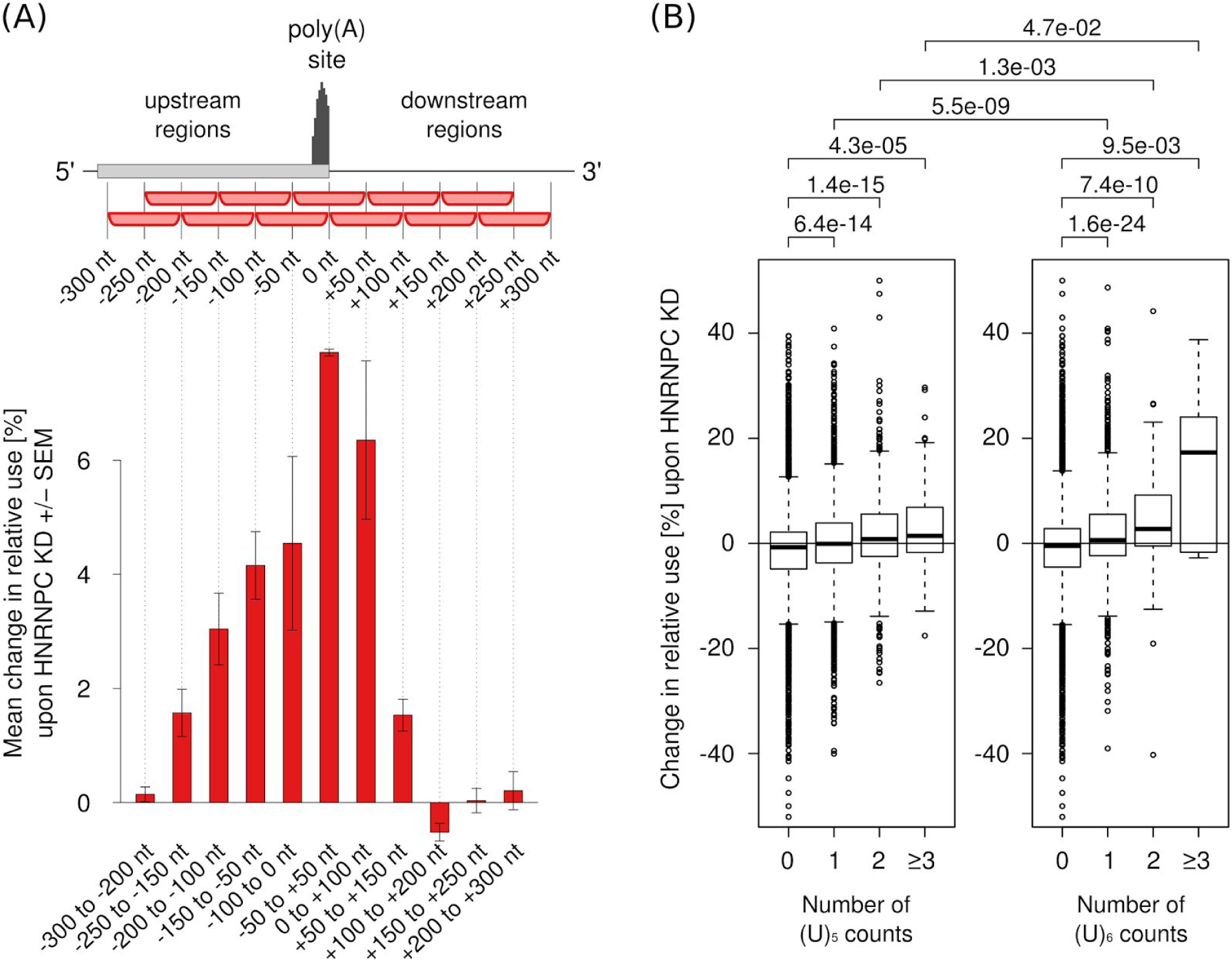
The length, number and location of poly(U) tracts with respect to poly(A) sites influence the change in poly(A) site use upon HNRNPC knock-down –. **(A)** Mean change in the use of sites containing the highest number of (U)_5_ motifs within 100 nucleotide long regions located at specific distances from the cleavage site (indicated on the x-axis) upon HNRNPC knock-down. Shown are mean +/- SEM in the two knock-down experiments. 250 poly(A) sites with the highest density of (U)_5_ motifs at each particular distance were considered. **(B)** Mean changes in the relative use of polyadenylation sites that have 0, 1, 2 or more (≥3) non-overlapping poly(U) tracts within +/- 50 nucleotides from their cleavage site. Distributions of relative changes in the usage of specific types of sites were compared and the p-values of the corresponding one-sided Mann-Whitney U tests are shown at the top of the panel.

Although the minimal RNA recognition motif of HNRNPC consists of five consecutive uridines (Ray et al. 2013; Liu et al. 2015; Cieniková et al. 2014), longer uridine tracts are bound with higher affinity (König et al. 2010; Cieniková et al. 2014; Zarnack et al. 2013). Consistently, we found that for a given length of the presumed HNRNPC binding site, the effect of the HNRNPC knock-down increases with the number of independent sites, and that given the number of non-overlapping poly(U) tracts, the effect of HNRNPC knock-down increases with the length of the sites (Figure 3B).

### Altered transcript regions contain ELAVL1 binding sites that mediate UDPL

As demonstrated above, binding of HNRNPC to U-rich elements that are located preferentially distally in terminal exons, seems to promote the use of proximal 3’ end processing sites. This leads to shorter 3’ UTRs, lacking U-rich regions with which a multitude of RNA binding proteins such as ELAVL1, (also known as Hu Antigen R, or HuR), could interact to regulate, among others, the stability of mRNAs in the cytoplasm (Brennan and Steitz 2001). To determine whether the HNRNPC-dependent alternative 3’ UTRs indeed interact with ELAVL1, we determined the number of ELAVL1 binding sites (obtained from a previous ELAVL1 crosslinking and immunoprecipitation study (Kishore et al. 2011)) that are located in the 3’ UTR regions between tandem poly(A) sites. To avoid any ambiguity, we used only terminal exons that contained precisely two poly(A) sites. As expected, we found a significant enrichment of ELAVL1 binding sites in 3’ UTR regions whose inclusion in transcripts changes in response to HNRNPC knock-down, compared to regions whose inclusion does not change (Figure 4A). Moreover, the density of ELAVL1 binding sites and not only their absolute number is enriched across these 3’ UTR regions (Figure 4B). Our results thus demonstrate that the HNRNPC-regulated 3' UTRs are bound, and probably susceptible to regulation by ELAVL1.

**Figure 4.**
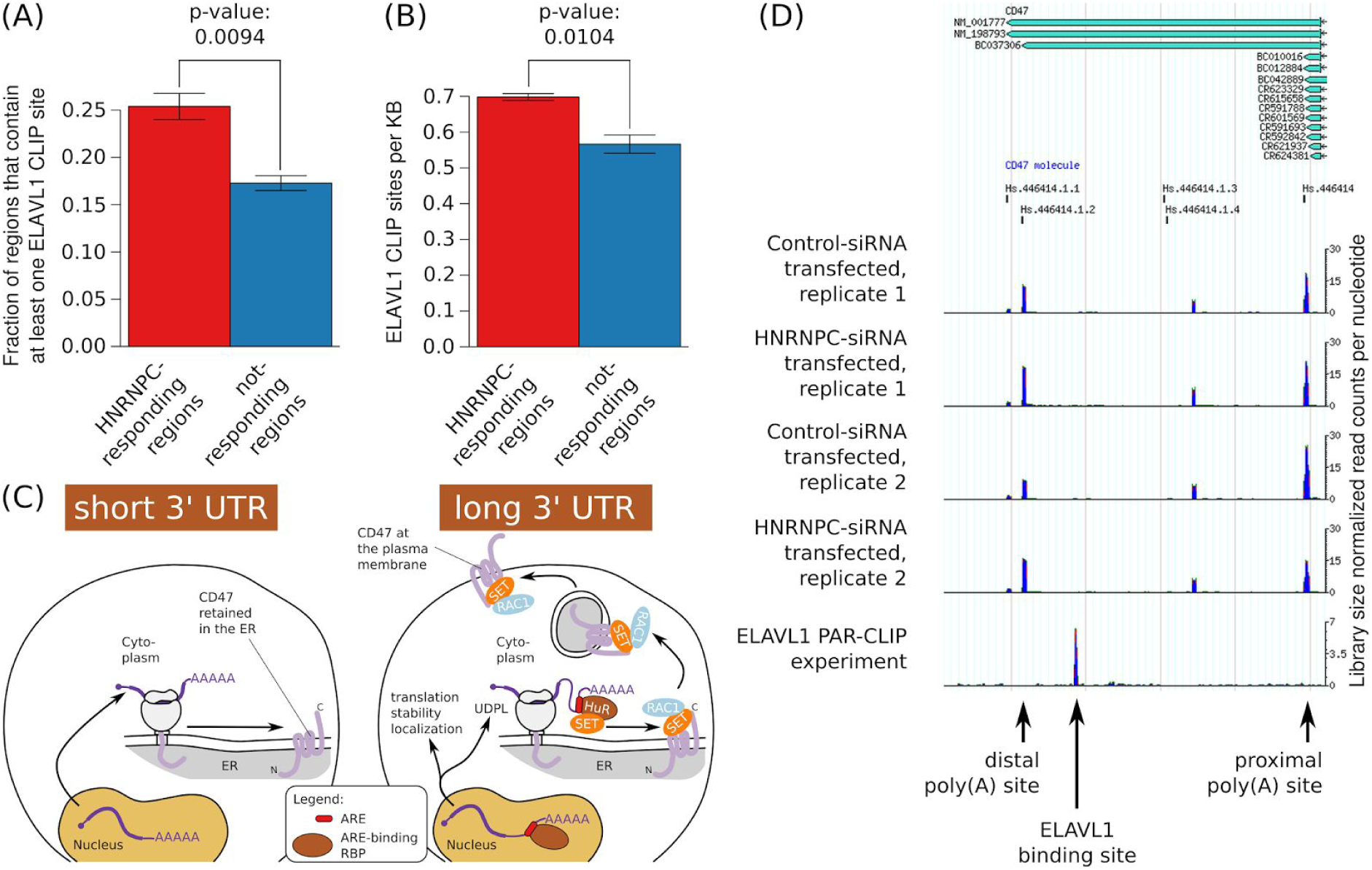
HNRNPC-responsive 3’ UTRs are enriched in ELAVL1 binding sites -. **(A)** Fraction of HNRNPC-responding and not-responding 3’ UTR regions that contain one or more ELAVL1 CLIP sites. The p-value of the one-sided t-test is shown. **(B)** Density of ELAVL1 CLIP sites (per kilobase) in the 3’ UTR regions described above. The p-value of the one-sided t-test is shown. **(C)** Model of the impact of A/U-rich elements (ARE) in 3’ UTR regions on various aspects of mRNA metabolism (Berkovits and Mayr 2015) **(D)** Density of A-seq2 reads along the CD47 3’ UTR in cells, showing the increased use of the distal poly(A) site in si-HNRNPC compared to si-Control transfected cells. The density of ELAVL1-CLIP reads in this region is also shown.

Recently, a new function has been attributed to the already multi-functional ELAVL1 protein. Work from the Mayr lab (Berkovits and Mayr 2015) showed that 3’ UTR regions that contain ELAVL1 binding sites can mediate 3’ UTR-dependent protein localization (UDPL). The ELAVL1 binding sites in the 3’ UTR of the CD47 molecule (CD47) transcript were found necessary and sufficient for the translocation of the CD47 transmembrane protein from the endoplasmatic reticulum to the plasma membrane, through the recruitment of the SET protein to the site of translation. SET binds to the cytoplasmic domains of the CD47 protein, translocating it from the endoplasmatic reticulum to the plasma membrane via active RAC1 (Berkovits and Mayr 2015; Klooster et al. 2007) (Figure 4C). Inspecting our data we found that the region of the CD47 3’ UTR that mediates UDPL is among those that responded to HNRNPC knock-down (Figure 4D). To find out whether HNRNPC can act as an upstream regulator of UDPL, we quantified the level of CD47 at the plasma membrane of cells that underwent siRNA-mediated knock-down of HNRNPC and cells that were treated with a control siRNA. Strikingly, we found that the CD47 level at the plasma membrane increased upon HNRNPC knock-down (Figures 5A and Supplementary Figure 10). Western blots for CD47 that were performed in HNRNPC and control siRNA-treated cells ruled out the possibility that the increase in membrane-associated CD47 upon HNRNPC knock-down was due to an increase in total CD47 levels (Supplementary Figure 11). We also carried out an independent immunofluorescence analysis of CD47 in these two conditions and again observed that the HNRNPC knock-down lead to an increase in the plasma membrane CD47 levels (Figure 5B). Overall, our results suggest that HNRNPC can function as an upstream regulator of UDPL.

**Figure 5.**
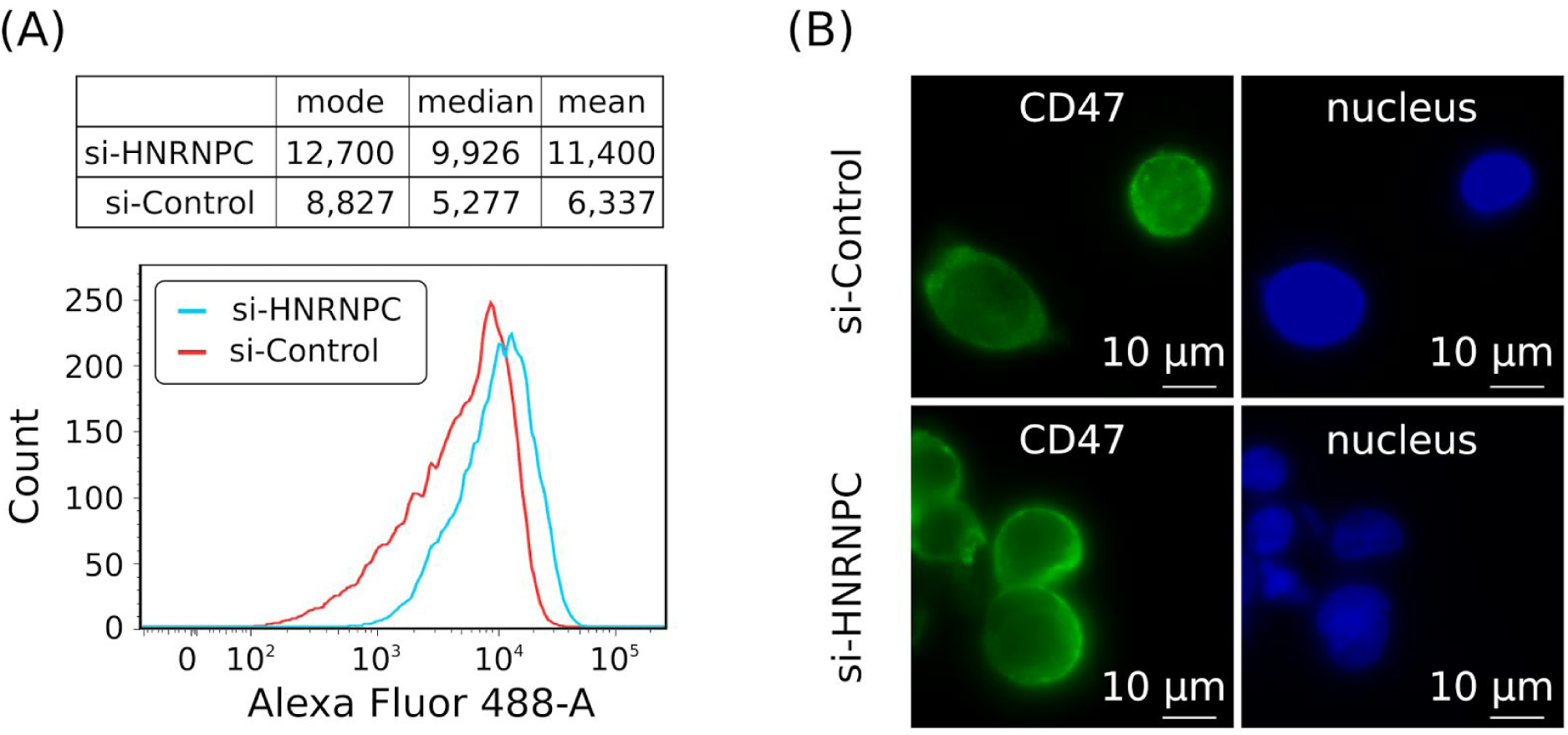
The knock-down of HNRNPC affects CD47 protein localization –. **(A)** Indirect immunophenotyping of membrane-associated CD47 in HEK 293 cells that were treated either with an si-HNRNPC (blue) or control (red) siRNA. Mean, median and mode of the Alexa Fluor 488 intensities computed for cells in each transfection set *(top panel)*, with histograms shown in the bottom panel. **(B)** Immunofluorescence staining of permeabilized HEK 293 cells with CD47 antibody (left panel) or nuclear staining with Hoechst (right panel). Top and bottom panels correspond to cells that were treated with control siRNA and si-HNRNPC, respectively.

### HNRNPC represses cleavage and polyadenylation at intronic, transcription start site-proximal poly(A) sites

Up to this point we focused on alternative polyadenylation (APA) sites that are located within single exons. However, given that HNRNPC binds to nascent transcripts, we also asked whether HNRNPC affects other types of APA, specifically at sites located in intronic regions. Indeed, we found that the HNRNPC knock-down increases the use of intronic poly(A) sites that are most enriched in putative HNRNPC-binding (U)_5_ motifs within +/- 50 nucleotides compared to sites that do not have (U)_5_ tracts within +/- 200 nucleotides (Figure 6A; p-values of the one-sided Mann-Whitney U test for the data from the two replicate knock-down experiments are: 1.4e-30 and 5.1e-29). To further validate the “masking” effect of HNRNPC on intronic poly(A) sites, we binned poly(A) sites into five groups based on their relative position within the host gene and asked how the position of sites within genes relates to their usage upon HNRNPC knock-down. As shown in Figure 6B, we found that intronic poly(A) sites that are most derepressed upon HNRNPC knock-down are preferentially located toward the 5’ ends of genes. We conclude that HNRNPC tends to repress the usage of intronic cleavage and polyadenylation sites whose usage leads to a strong reduction of transcript length. Figure 6C shows the example of the Kelch-Like Family Member 3 (KLHL3) gene, which harbors one of the most derepressed intronic poly(A) sites.

**Figure 6.**
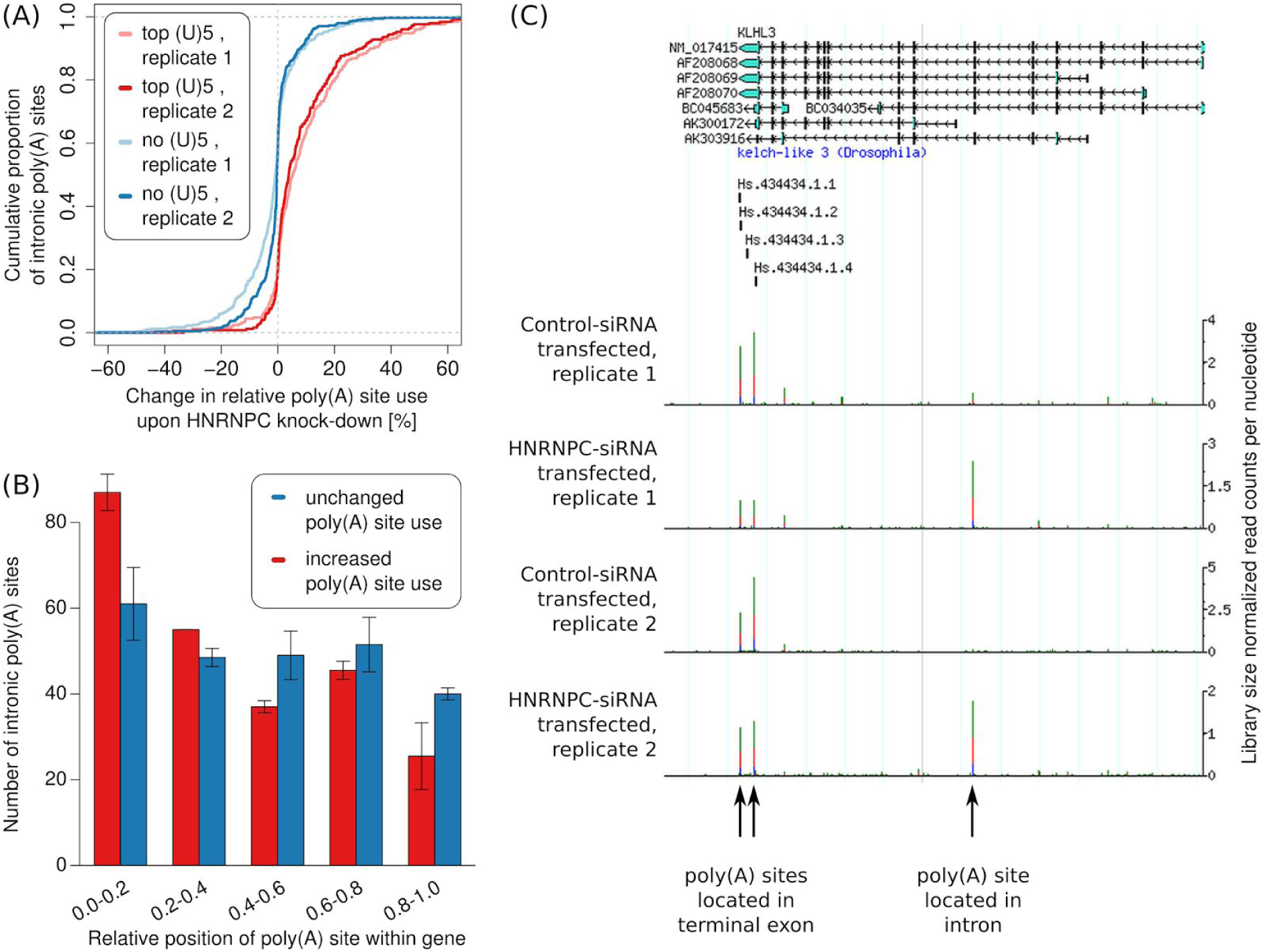
- HNRNPC knock-down leads to increased usage of intronic poly(A) sites. **(A)** The change in the relative use of intronic poly(A) sites that do not contain any (U)_5_ within +/- 200 nucleotides and of the top 250 intronic poly(A) sites according to the number of (U)_5_ motifs within +/- 50 nucleotides around the cleavage site, upon HNRNPC knock-down. **(B)** Relative location of the top 250 most de-repressed intronic poly(A) sites that have HNRNPC binding motifs within -200 to +100 nts around their cleavage site and of the 250 intronic poly(A) sites that changed least upon HNRNPC knock-down. **(C)** Screenshot of the KLHL3 gene, in which intronic cleavage and polyadenylation is strongly increased upon HNRNPC knock-down.

## Discussion

Studies in recent years have shown that pre-mRNA cleavage and polyadenylation is a dynamically regulated process that yields transcript isoforms with distinct interaction partners, subcellular localization, stability and translation rate (for a review see e.g. Davis and Shi 2014). Specific polyadenylation programs seem to have evolved in relation with particular cell types or states. For example, APA and 3’ UTR lengths are developmentally regulated (Ji et al. 2009; Ji and Tian 2009; Miura et al. 2013) and short 3’ UTRs are generated in proliferating and malignant cells (Lee et al. 2007; Sandberg et al. 2008; Xia et al. 2014). The key regulators of these polyadenylation programs are unknown. Reduced expression of the U1 snRNP (Berg et al. 2012) or of the mammalian cleavage factor I (CFI_m_) components NUDT21 and CPSF6 (Martin et al. 2012; Gruber et al. 2012) can cause a systematic reduction in 3’ UTR lengths, but only limited evidence about the relevance of these factors in physiological conditions has been provided (Berg et al. 2012; Masamha et al. 2014). Other factors that are part of the 3’ end processing machinery and have systematic effects on polyadenylation are the nuclear poly(A) binding protein 1 (Jenal et al. 2012), which suppresses cleavage and polyadenylation, the 64 kDa cleavage stimulation factor (CSTF2) component of the 3’ end cleavage and polyadenylation complex whose expression correlates with the preferential use of short 3’ UTRs in cancer cells (Xia et al. 2014) and the retinoblastoma binding protein 6, whose reduced expression results in reduced transcript levels and increased use of distal poly(A) sites (Di Giammartino et al. 2014).

Many experimental protocols to capture transcript 3’ ends and enable studies of the dynamics of polyadenylation have been developed (reviewed in de Klerk et al. 2014), and consequently, a few databases of 3’ end processing sites are available (Lee et al. 2007; Derti et al. 2012; You et al. 2014). However, none of these databases has used the entire set of 3’ end sequencing data available today and thus their coverage is limited. In this study, we have developed a procedure to automatically process heterogeneous data sets generated with one of nine different protocols, aiming to identify *bona fide* poly(A) sites that are independently regulated. Although most of the reads that were used to construct the currently available databases (Derti et al. 2012; You et al. 2014) map within the poly(A) site clusters that we constructed, the differences at the level of reported processing sites are quite large. This is largely due to the presence of many sites with very limited read support and no upstream poly(A) signals in previous data sets. For example, focusing on the terminal exons of protein-coding genes and lincRNAs from the UCSC Basic table of the GENCODE V19 annotation, the human atlas that we constructed has a higher fraction of exons with assigned poly(A) sites compared to previous databases. 71.12 % of all terminal exons of protein coding genes in our atlas have at least one annotated poly(A) site in contrast to 66.26 % and 62.69 % for the studies of Derti et al. (Derti et al. 2012) and You et al. (You et al. 2014), respectively. The coverage of the terminal exons of lincRNAs is smaller overall, but again clearly higher in our atlas (37.59%) compared to those of Derti et al. (Derti et al. 2012) and You et al. (You et al. 2014) (29.57% and 24.51%, respectively, Supplementary Figure 12). The lower coverage of lincRNAs is probably due to their lower expression in comparison with protein-coding genes (Wu et al. 2014) and to the fact that some of them are bimorphic, appearing in both the poly(A)^+^ and poly(A)" fraction, or not polyadenylated (Hangauer et al. 2013) and thereby cannot be captured efficiently with protocols that require the presence of a poly(A) tail.

Although for mouse we did not have lincRNA annotations, the general trend of higher coverage in our atlas compared to existing ones holds for mouse genes as well (Supplementary Figure 13; detailed numbers in Supplementary Tables 7 and 8).

The 3’ end processing sites reported by other studies (Derti et al. 2012; You et al. 2014) but missing from our atlas have, on average, a substantially lower read support. Some were only documented by multi-mapping reads, had features indicative of internal priming, or originated in regions from which broadly scattered reads were generated.

Building upon a large set of 3’ end sequencing samples, we have analyzed the sequence composition around high-confidence poly(A) sites to identify elements that may recruit RNA-binding proteins to modulate polyadenylation. We have identified sequence motifs that exhibit a positional preference with respect to 3’ end cleavage sites almost identical to the canonical poly(A) signal AAUAAA. Six of the ten novel motifs that we found in each human and mouse data set are shared. With a more comprehensive set of poly(A) signals we have been able to more efficiently use data from many heterogeneous experiments, thereby achieving a higher coverage of terminal exons and annotated genes by poly(A) sites. Even though the poly(A) and poly(U) motifs are also strongly enriched around poly(A) sites, they were not annotated as poly(A) signals due to positional profiles divergent from what is expected for poly(A) signals. The general A- and U-richness of the neighborhood of cleavage and polyadenylation sites has been observed before (Tian and Graber 2012), but the RNA-binding protein interactors and their role in polyadenylation remain to be characterized.

Here we hypothesized that HNRNPC, a protein that binds poly(U) tracts (Cieniková et al. 2014; Liu et al. 2015; Ray et al. 2013) and has a variety of functions including pre-mRNA splicing (König et al. 2010) and mRNA transport (McCloskey et al. 2012), also modulates the processing of pre-mRNA 3’ ends. HNRNPC has originally been identified as a component of the HNRNP core particle (Beyer et al. 1977; Choi and Dreyfuss 1984) and found to form stable tetramers that bind to nascent RNAs (Whitson et al. 2005). Systematic evolution of ligands by exponential enrichment (SELEX) experiments have shown that HNRNPC particles bind to uninterrupted tracts of five or more uridines (Görlach et al. 1994), and studies employing crosslinking and immunoprecipitation indicated that longer tracts are bound with higher affinity (König et al. 2010). By sequencing mRNA 3’ ends following the siRNA-mediated knock-down of HNRNPC we found that transcripts that contain poly(U) tracts around their poly(A) sites respond in a manner indicative of HNRNPC masking poly(A) sites. This is reminiscent of the U1 snRNP protecting nascent RNAs from premature cleavage and polyadenylation, in a mechanism that has been called “telescripting” (Kaida et al. 2010; Berg et al. 2012). Indeed, HNRNPC seems to have at least in part a similar function, because the knock-down of HNRNPC increases the incidence of cleavage and polyadenylation at intronic sites, with a preference for intronic sites close to the transcription start. It should be noted that these intronic sites are not spurious, but have experimental support as well as polyadenylation signals. Thus, the short transcripts that terminate at these sites could be functionally relevant, either through the production of truncated proteins or through an effective down-regulation of the functional, full-length transcript forms. In terminal exons, U-rich poly(A) sites whose usage increases upon HNRNPC knock-down tend to be located distally. In these transcripts, HNRNPC may function to “mask” the distal, “stronger” signals, allowing the “weaker” proximal poly(A) sites to be used (Shi 2012). Interestingly, the competition between HNRNPC and U2AF2 appears to prevent exonization of Alu elements (Zarnack et al. 2013) and furthermore, polyadenylation at Alu elements is regulated by HNRNPC (Tajnik et al. 2015). These studies have emphasized the complex cross-talks between regulators that come into play during RNA splicing and polyadenylation (Proudfoot 2011). They also illustrate the striking multi-functionality of U-rich and A/U-rich elements that are bound by various proteins at different stages to modulate processes ranging from transcription termination (Almada et al. 2013) up to protein localization (Berkovits and Mayr 2015).

Initial studies that reported 3’ UTR shortening in dividing cells hypothesized that shortened 3’ UTRs harbor a reduced number of miRNA binding sites, the corresponding mRNAs being more stable and having an increased translation rate (Sandberg et al. 2008; Mayr and Bartel 2009). However, genome-wide measurements of mRNA and protein levels in dividing and resting cells revealed that systematic 3’ UTR shortening has a relatively minor impact on mRNA stability, translation and protein output (Gruber et al. 2014; Spies et al. 2013). Instead, evidence has started to emerge that 3’ UTR shortening results in the loss of interaction with various RNA-binding proteins, whose effects are not limited to mRNA stability and translation (Gupta et al. 2014), but reach as far as the transport of transmembrane proteins to the cytoplasmic membrane (Berkovits and Mayr 2015). The CD47 protein provides a striking example of 3’ UTR-dependent protein localization. However, the upstream signals and perhaps additional targets of this mechanism remain to be uncovered. Here we have demonstrated that HNRNPC can modulate polyadenylation of a large number of transcripts, leading to the inclusion/removal of U-rich elements. When these elements remain part of the 3’ UTRs, they can be subsequently bound by a variety of U-rich element binding proteins, including ELAVL1, which has been recently demonstrated to play a decisive role in the UDPL of CD47 (Berkovits and Mayr 2015). Indeed, we found that the knock-down of HNRNPC promotes the expression of the long CD47 3’ UTR that is accompanied by an increased membrane localization of the CD47 protein. Although HNRNPC does not appear to target any particular class of transcripts, nearly one quarter (≥23%) of the HNRNPC-responsive transcripts encode proteins that are annotated with the Gene Ontology category "integral component of membrane" (GO:0016021). Thus, our results provides an extended set of candidates for the recently discovered UDPL mechanism.

In conclusion, PolyAsite, available at http://polyasite.unibas.ch is a large and extendable resource that supports investigations into the polyadenylation programs that operate during changes in cell physiology, during development and in malignancies.

## Methods

### Uniform processing of publicly available 3' end sequencing data sets

Publicly available 3' end sequencing data sets were obtained from the NCBI GEO archive (www.ncbi.nlm.nih.gov/geo) and from NCBI SRA (www.ncbi.nlm.nih.gov/sra). To ensure uniform processing of 3' end sequencing data generated by diverse 3' end sequencing protocols, we developed the following computational pipeline (Supplementary Figure 14). First, raw sequencing files were converted to FASTA format. For samples generated with protocols that leave a 5' adapter sequence in the reads, we only retained the reads from which the specified adapter sequence could be trimmed. Next, we trimmed the 3' adapter sequence and, when the protocol captured the reverse complement of the RNAs, we reverse complemented the reads. Reads were then mapped to the corresponding genome assembly (hg19 and mm10, respectively) and to mRNA and lincRNA-annotated transcripts (GENCODE 14 release for human and ENSEMBL annotation of mouse, both obtained from UCSC June 2013). The sequence alignment was done with segemehl version 0.1.4–395 with default parameters (Hoffmann et al. 2009). In cases where the sex of the organism from which the sample was prepared was female, mappings to the Y chromosome were excluded from further analysis. For each read, we only kept the mappings with the highest score (smallest edit distance). Mappings to the transcriptome were only considered if they overlapped a splice junction with at least 5 nt on both sides and they had a higher score compared to any mapping of the same read to the genomic sequence. Based on the genome coordinates of individual exons and the mapping coordinates of reads within transcripts, we next converted read-to-transcript mapping coordinates into read-to-genome mapping coordinates. For generating a high confidence set of pre-mRNA 3' ends we started from reads that consisted of no more than 80 % of adenines and that mapped uniquely to the genome such that the last three nucleotides of the read were perfectly aligned. Furthermore, we required that the 3' end of the read was not an adenine. Finally, we collapsed the 3' ends of the sequencing reads into putative 3' end processing sites and filtered out those that likely originated from internal priming (for details on the entire pipeline see Supplementary Figure 14).

### Clustering of closely spaced 3' end sites into 3' end processing regions

Putative 3' end processing sites identified as described above were used to construct clusters to (1) identify poly(A) signals, (2) derive sample-specific cutoffs for the number of reads necessary to support a site and (3) determine high-confidence 3' end processing sites in the human and mouse genome. In clustering putative 3' end processing sites from multiple samples, as done for analyses (1) and (3), we first sorted the list of 3' end sites by the number of supporting samples and then by the total normalized read count (read counts were normalized per sample as reads per million (RPM) and for each site a total RPM was obtained by summing these numbers over all samples). In contrast, to generate clusters of putative reads from individual samples (analysis 2.), we only ranked genomic positions by RPM. Clusters were generated by traversing the sorted list from top to bottom and associating lower-ranking sites with a representative site of a higher rank, if the lower-ranked sites were located within a specific maximum distance upstream d_u_ or downstream d_d_ of the representative site (Supplementary Figure 15).

To determine a maximum distance between sites that seem to be under the same regulatory control, we applied the above-described clustering procedure for distances d_u_ and d_d_ varying between 0 and 25 nucleotides and evaluated how increasing the cluster length affects the number of generated clusters that contain more than one site (Supplementary Figure 16). Consistent with previous observations, we found at a distance of 8 nucleotides from the representative site ~40 % of the putative 3' end processing sites are part of multi-site clusters, this proportion increases to 43 % for a distance of 12 nts and reaches 47 % at a distance of 25 nts. For consistency with previous studies, we used d_u_ = d_d_ = 12 nt (Tian et al. 2005; You et al. 2014). Only for the clustering of putative 3' end processing sites in individual samples we used a larger distance, d_u_ = d_d_ = 25, resulting in a more conservative set of clusters, with a maximum span of 51 nucleotides.

### Identification of poly(A) signals

To obtain a set of high-confidence 3' end processing sites from which to identify poly(A) signals, we filtered the preliminary 3' end clusters, retaining only those that were supported by data from at least two protocols. For clusters with at least two putative sites we took the center of the cluster as the representative cleavage site. Then, we constructed the positional frequency profile in the [-60,-5] nucleotides region upstream of the representative cleavage sites for each of the 4096 possible hexamers (Supplementary Figure 17A). We did not consider the five nucleotides upstream of the putative cleavage sites, to reduce the impact of artifacts originating from internal priming at poly(A) nucleotides, which are very close in sequence to the main poly(A) signal, AAUAAA (see “Treatment of putative 3' end sites originating from internal priming”). Before fitting a specific functional form to the frequency profiles we smoothed them, taking at each position the average frequency in a window of 11 nucleotides centered on that position, and we subtracted a motif-specific "background" frequency which we defined as the median of the ten smallest frequencies of the motif in the entire 55 nucleotides window. To identify motifs that have a specific positional preference upstream of the cleavage site, we fitted a Gaussian density curve to the background-corrected frequency profile with the "nls" function in R, assessing the quality of the fit by the r^2^ value and by the height:width ratio of the fitted peak, whereas the width was defined as the standard deviation of the fitted Gaussian density (Supplementary Figure 17A). Alternative poly(A) signals should have the same positional preference as the main signal, AAUAAA. However, when considering 60 nts upstream of the cleavage site, poly(A) signals can occur not only at -21 nts, which seems to be the preferred location of these signals, but also at other positions, particularly when the poly(A) signal is suboptimal and co-occurs with the main signal. Thus, we started from motifs that peaked in the region upstream of the cleavage site (r^2^ ≥ 0.6 for the fit to the Gaussian and a height:width ratio ≥ 5), but allow a permissive position of the peak, between -40 to -10 nts. Putative poly(A) signals were then determined according to the following iterative procedure (Supplementary Figure 17B):

We sorted the set of putative signals by their strength. The strongest signal was considered to be the one with the lowest p-value of the test that the peak frequency of the motif could have been generated by Poisson sampling from the background rate inferred as the mean motif frequency in the regions of 100 to 200 nts up- and downstream of the cleavage site. As expected, in both human and mouse data sets the most significant hexamer was the canonical poly(A) signal AAUAAA. At every iteration, we removed all sequences that contained the most significant signal in the -60 nts window upstream of the cleavage sites and repeated the procedure on the remaining set of sequences. Signals with an r^2^ value of the fit to a Gaussian ≥ 0.9 and a height:width ratio ≥ 4 were retained, the most significant retained and the procedure was iterated. The fitted Gaussian densities of almost all of the putative poly(A) signals recovered with this procedure had highly similar peak positions and standard deviations. Therefore, only signals that peaked at most one nt away from the most significant hexamer, AAUAAA, were retained in the final set of poly(A) signals. The only hexamers that did not satisfy this condition were the AAAAAA hexamer in mouse and AAAAAA as well as TTAAAA in human.

### Treatment of putative 3' end sites originating from internal priming

Priming within A-rich, transcript-internal regions rather than to the poly(A) tail is known to lead to many false positive sites with most of the existing 3' end sequencing protocols. We tried to identify and eliminate these cases as described in section “Uniform processing of publicly available 3' end sequencing data sets”. An under-appreciated source of false positives seems to be the annealing of the poly(T) primer in the region of the poly(A) signal itself, which is A-rich and close to the poly(A) site (Tian et al. 2005; Shi 2012). Indeed, a preliminary inspection of cleavage sites that seemed to lack poly(A) signals revealed that these sites were located on or in the immediate vicinity of a motif that could function as a poly(A) signal. To reduce the rate of false positives generated by this mechanism we undertook an additional filtering procedure as follows (Supplementary Figure 18). First, every 3' end site that was located within a poly(A) signal or had a poly(A) signal starting within five nucleotides downstream of the apparent cleavage site was marked initially as "PAS priming site". Then, during the clustering procedure, each cluster that contained a "PAS priming site" was itself marked as putative internal priming candidate and the most downstream position of the cluster was considered as the representative site for the cluster. Finally, internal priming candidate clusters were either (i) merged into a downstream cluster, if all annotated poly(A) signals of the downstream cluster were also annotated for the internal priming candidate or (ii) retained as valid poly(A) cluster when the distance between the representative site to the closest poly(A) signal upstream was at least 15 nts or (iii) discarded, if neither condition (i) nor (ii) were met.

### Generation of the comprehensive catalog of high-confidence poly(A) sites

#### Annotating poly(A) signals

The procedure outlined in the sections above yielded 18 signals that showed a positional preference similar to AAUAAA in both mouse and human. These signals were used to construct the catalog of 3' end processing sites. We started again from all unique apparent cleavage sites from the 78 human and 110 mouse samples (Supplementary Tables 3 and 4), amounting to 6,983,499 and 8,376,450 sites, respectively. For each of these sites, we annotated all occurrences of any of the 18 poly(A) signals within -60 to +5 nts relative to the apparent cleavage site.

#### Identification of 3' end processing clusters expressed above background in individual samples

For each sample independently, we constructed clusters of 3' end processing sites as described above (see “Clustering of closely spaced 3' end sites into 3' end processing regions”). At this stage, we did not eliminate "PAS priming sites", but rather used a larger clustering distance, of d_u_ = d_d_ = 25 (see “Clustering of closely spaced 3' end sites into 3' end processing regions”), to ensure that "PAS priming sites" were captured as well. We kept track of whether any 3' end processing site in each cluster had an annotated poly(A) signal or not. Next, we sorted the clusters by the total number of reads that they contained and, traversing the sorted list from top (clusters with most reads) to bottom, we determined the read count *c* at which the percentage of clusters having at least one annotated poly(A) signal, dropped below 90%. We then discarded all clusters with ≤ *c* read counts, as not having sufficient experimental support (Supplementary Figure 19 outlines how to determine sample-specific cutoffs).

#### Combining poly(A) site clusters from all samples into a comprehensive catalog of 3' end processing sites

Starting from the sites identified in at least one of the samples we first normalized the read counts to the total number of reads in each sample to compute expression values as reads per million (RPM), then merged all sites into a unique list which we sorted first by the number of protocols supporting each individual site and then by the total RPM across all samples that supported the site. These sites were clustered as described above in the section “Clustering of closely spaced 3' end sites into 3' end processing regions” and then internal priming candidates were eliminated as described in section “Treatment of putative 3' end sites originating from internal priming”. Closely spaced clusters were merged: (i) when they shared the same poly(A) signals, or (ii) when the length of the resulting cluster did not exceed 25 nucleotides. The above procedure could result in poly(A) clusters that were still close to each other but with a combined length exceeding the maximum cluster size, and that did not have any poly(A) signal annotated. To retain from these the most likely and distinct poly(A) sites we merged clusters without poly(A) signals with an inter-cluster-distance ≤ 12 nts and retained those whose total cluster span was ≤ 50 nucleotides. A small fraction of the clusters had a span ≥ 50 nucleotides with some even wider than 100 nts. These clusters were not included in the atlas. Finally, the position with the highest number of supporting reads in each cluster was reported as the representative site of the cluster (Supplementary Figure 20). The final set of clusters was saved in a BED-formatted file, with the number of supporting protocols as the cluster score. A cluster obtained support by a protocol if any of the reads in the clusters originated from that protocol. We used the mRNA and lincRNA annotations from the UCSC Basic table of the GENCODE 19 version for human and the ENSEMBL mm10 transcript annotation from UCSC for mouse to annotate the following categories of clusters, listed here in the order of their priority (which we used to resolve annotation ambiguity):

TE: terminal exon
EX: any other exon except the terminal one
IN: any intron
DS: up to 1000 nt downstream of an annotated gene
AE: antisense to an annotated exon
AI: antisense to an annotated intron
AU: antisense and within 1000 nucleotides upstream of an annotated gene
IG: intergenic.

#### Sequence logos of the identified poly(A) signals

The procedure described above (see “Combining poly(A) site clusters from all samples into a comprehensive catalog of 3' end processing sites”) was used to construct a version of the human and mouse poly(A) site atlases that incorporated the entire set of 22 organism-specific poly(A) signals, not just the 18 signals that were shared between species. Frequencies of all annotated poly(A) signals (possibly more than one per poly(A) cluster) across all identified clusters were calculated for the human and mouse catalog independently. Fasta files with poly(A) signals, including their multiplicities in the data were used with the Weblogo program (Crooks et al. 2004) version 3.3, with default settings, to generate the sequence logos for human and mouse, respectively.

#### Hexamer enrichment in upstream regions of 3' end clusters

We calculated the significance (p-value) of enrichment of each hexamer in the set of 3' end clusters (and their 60 nt upstream regions) of our human and mouse atlas relative to what would be expected by chance, assuming the mono-nucleotide frequencies of the sequences and a binomial distribution of motif counts.

#### Annotation of poly(A) sites with respect to categories of genomic regions

We used the genomic coordinates of the protein-coding genes and lincRNAs from the UCSC Basic table of the GENCODE V19 (human) and the ENSEMBL mm10 (mouse) annotations to annotate ours and previously published sets of poly(A) sites with respect to genomic regions with which they overlap. A poly(A) site was assigned to an annotated feature if at least one of its genomic coordinates overlapped with the genomic coordinates of the feature.

**PolyAsite:** For every poly(A) cluster annotated in our catalogs, the entire region of the cluster was used to test for an overlap with annotated genomic features.

**PolyA-seq:** Processed, tissue-specific data were downloaded as BED-file (http://www.ncbi.nlm.nih.gov/geo/query/acc.cgi?acc=GSE30198). Poly(A) sites from nine and five different samples were downloaded for human and mouse, respectively (Derti et al. 2012). Mouse genome coordinates were converted to the coordinates of the ENSEMBL mm10 annotation through LiftOver (Hinrichs et al. 2006). The genomic coordinates of all poly(A) sites (one position per poly(A) site) were intersected with the annotation features.

**APASdb:** Processed, tissue-specific data for human poly(A) sites were downloaded from http://mosas.sysu.edu.cn/utr/download_datasets.php. This included poly(A) sites from 22 human tissues (You et al. 2014). The genomic coordinates of all poly(A) sites (one position per poly(A) site) were intersected with the annotation features.

### Analysis of 3' end libraries from HNRNPC knock-down experiments

#### Sequencing of A-seq2 libraries and quantification of relative poly(A) site usage

We considered all high-confidence A-seq2 (Gruber et al. 2014) reads that mapped to a unique position in the human genome (hg19) and whose 5’ ends were located in a cluster supported by two or more protocols. For our A-seq2 protocol, high-confidence reads are defined as sequencing reads that do not contain more than 2 ambiguous bases (N), have a maximum A-content of 80% and the last nucleotide is not an adenine. Using our atlas of poly(A) sites that was constructed considering the 18 conserved poly(A) signals, we calculated the relative usage of poly(A) sites. Because we focused mostly on tandem poly(A) sites we considered in our analysis all exons that had multiple poly(A) clusters expressed at more than 3.0 RPM in one or more samples. There were 12,136 such clusters. As “consistent” changing poly(A) sites we considered those that had a change of at least 5% in the same direction in both replicates. And as “consistent“ minor/non-changing poly(A) sites we considered sites whose mean change and standard deviation across replicates were less than 2%.

#### Determination of ELAVL1 binding sites that are affected by APA events taking place upon HNRNPC knock-down

##### Determination of 3' UTR regions that respond to HNRNPC knock-down

To identify putative HNRNPC regulated regions we have selected exons that had exactly two poly(A) sites, one of which showing an increase in relative usage by at least 5% upon HNRNPC knock-down and harboring a putative HNRNPC binding site ((U)_5_) within a region of -200 to 100 nucleotides relative to the cleavage site. As minor/non-changing regions we considered exons having exactly two poly(A) sites, both of which are changing less than 5% upon HNRNPC knock-down.

##### ELAVL1 binding site extraction from CLIP experiments

We used data from a previously published ELAVL1 CLIP experiment (Kishore et al. 2011), Gene Expression Omnibus (GEO, http://www.ncbi.nlm.nih.gov/geo/) database accession GSM714641. Enriched binding sites were determined by applying the mRNA site extraction tool available on CLIPZ (Khorshid et al. 2011; Jaskiewicz et al. 2012) using the mRNA-Seq samples with GEO accessions GSM714684 and GSM714685 as background. CLIP sites with an enrichment score ≥ 5.0 were translated into genome coordinates (hg19) using GMAP (Wu and Watanabe 2005). To identify ELAVL1 CLIP sites located within transcript regions that are included/excluded through alternative polyadenylation, we intersected the set of enriched ELAVL1 CLIP sites with genomic regions enclosed by tandem poly(A) sites (located on the same exon) using BEDtools (Quinlan and Hall 2010). To evaluate the relative positioning of ELAVL1 binding sites within regions between tandem poly(A) sites, the relative positions of the binding sites were calculated using the center of each CLIP site as its coordinate.

### Determination of intronic poiy(A) sites

To make sure that we can capture premature cleavage and polyadenylation events that might occur spontaneously upon knock-down of HNRNPC and are therefore observable in the HNRNPC knock-down samples only, for each sample we created clusters as described above (see “Clustering of closely spaced 3' end sites into 3' end processing regions”), using conserved poly(A) signals only. By analogy to tandem poly(A) sites within exons, we calculated the relative usage of clusters within genes by considering all genes having multiple poly(A) clusters that were expressed at greater than 3.0 RPM in one or more samples. There were 22,498 such clusters. Finally, we determined the set of sites that showed a consistent change upon HNRNPC knockdown as described above (see “Sequencing of A-seq2 libraries and quantification of relative poly(A) site usage”).

### Cell culture and RNAi

HEK 293 cells (Flp-In™-293, from Life Technologies) were grown in Dulbecco’s modified Eagle's medium (DMEM, from Sigma) supplemented with 2 mM L-glutamine (GIBCO) and 10% heat-inactivated fetal calf serum (GIBCO). Transfections of siRNA were carried out using Lipofectamine RNAiMAX (Life Technologies) following manufacturer’s protocol. The following siRNAs were used. Negative-control from Microsynth (sense strand AGG UAG UGU AUC GCC UUG TT), and si-HNRNPC1/2 (sc-35577 from Santa Cruz Biotechnologies), both applied at 20 nM in 2.5 ml DMEM on 6-well plates.

### Western blotting

Cells were lysed in 1 x RIPA Buffer and protein concentration was quantified using BCA reagent (Thermo Scientific). A stipulated amount of the sample (usually 10 microgram) was then used for SDS gel separation and transferred to ECL membrane (Protran, GE Healthcare) for further analysis. Membranes were blocked in 5\% skim milk (Migros) in TN-Tween (20 mM Tris-Cl pH 7.5, 150 mM NaCl, 0.05% Tween-20). The following antibodies were used for Western blots. Actin, sc-1615 from Santa Cruz Biotechnology; hnRNP-C, sc-10037 from Santa Cruz Biotechnology; CD47, AF-4670 from R&D Systems. HRP-conjugated secondary antibodies were applied at 1:2000 dilution. After signal activation with ECL Western blotting detection reagent (GE Healthcare), imaging of Western blots was performed on an Azure c600 system. Signal quantification was done with ImageJ software.

### Immunofluorescence

For the immunofluorescence analysis, HEK 293 cells were transfected with either control siRNA or siRNAs targeting HNRNPC as described under Cell culture and RNAi, 48 hours post transfection cells were fixed with 4% paraformaldehyde for 30 min, permeabilized and blocked with PBS containing 1% BSA and 0.1% Triton X-100 for 30 min. Primary anti-CD47 antibody (sc-59079 from Santa Cruz Biotechnology) was incubated for 2 h at room temperature at a dilution of 1:100 in the same buffer. To visualize CD47 in cells, secondary antibody conjugated with Alexa Fluor 488 was applied, while the nucleus was labelled with Hoechst dye. Imaging was performed with a Nikon Ti-E inverted microscope adapted with a LWD condenser (WD 30mm; NA 0.52), Lumencor SpectraX light engine for fluorescence excitation LED transmitted light source. Cells were visualized with a CFI Plan Apochromat DM 60X Lambda oil (NA 1.4) objective and images were captured with a Hamatsu Orca-Flash 4.0 CMOS camera. Image analysis and edge detection was performed with NIKON NIS Elements software Version 4.0. All images were subsequently adjusted uniformly and cropped using Adobe Photoshop CS5.

### FACS Analysis

FACS analyses of siRNA transfected cells were performed similar to immunofluorescence studies (see “Immunofluorescence”) except that cells were not permeabilized prior to the treatment with antibody against CD47 (sc-59079 from Santa Cruz Biotechnology). Analysis of Alexa Fluor 488 signal and counts was carried out on a BD FACS Canto II instrument and data was analyzed with the FLOWJO software. An equal pool of siRNA samples from each transfection set was mixed for the IgG control staining to rule out nonspecific signals.

### PAR-CLIP and A-seq2 libraries

A-seq2 libraries were generated as described (Gruber et al. 2014) and sequenced on a Illumina HiSeq 2500 deep sequencer. The HNRNPC PAR-CLIP was performed as described (Martin et al. 2012) with a modification consisting of preblocking of the Dynabeads-protein-A (Life Technologies) resulting in reduced background and higher efficiency of library generation. To this end, Dynabeads were washed 3x with PN8 buffer (PBS buffer with 0.01% NP-40), incubated in 0.5 ml of PN8-preblock (1 mM EDTA, 0.1% BSA from Sigma, A9647, and 0.1 mg/ml heparin, from Sigma, H3393, in PN8 buffer) for 1 hr on a rotating wheel. The preblock solution was removed and replaced by the antibody in 0.2 ml preblock solution and rotated for 2–4 hrs. We used the goat polyclonal antibody sc-10037 against HNRNPC1/C2 (Santa Cruz Biotechnology). Used 5' adapter: GTT CAG AGT TCT ACA GTC CGA CGA TC and 3' adapter: TGG AAT TCT CGG GTG CCA AGG.

### HNRNPC PAR-CLIP analysis

The raw data were mapped using CLIPZ (Khorshid et al. 2011). For each poly(A) site the uniquely mapping reads that overlapped with a region of +/-50 nts around the cleavage site were counted and normalized (divided) by the expression level (RPKM) of the poly(A) sites host gene using the mRNA-Seq samples with GEO accessions GSM714684. For Supplementary Figure 9 normalized CLIP read counts of poly(A) sites belonging to different categories of consistently behaving poly(A) sites across replicates, as defined above (see “Sequencing of A-seq2 libraries and quantification of relative poly(A) site usage”), were used.

## Data access

The sequencing data have been submitted to the sequence read archive (SRA) of the National Center for Biotechnology Information with the accession number SRP065825.

## Acknowledgements

This work was supported by the Swiss National Science Foundation grant #31003A-143977 to W.K. and by the Swiss National Science Foundation NCCR project “RNA & Disease”.

We would like to thank Erik van Nimwegen for his input on data analysis, Béatrice Dimitriades for technical assistance and Josef Pasulka for suggestions on the analysis.

## Author contributions

M.Z., A.J.G., A.R.G. and W.K. have designed the project. A.R.G., R.S. and A.J.G. have collected datasets and created the catalog of poly(A) sites. M.B. has developed the PolyAsite web interface. R.S. and A.J.G. have identified poly(A) signals. G.M. has performed the HNRNPC PAR-CLIP and A-seq2 experiments. A.J.G. has analyzed the data with help from R.S. G.M. and S.G. have performed the experiments on CD47. M.Z. has supervised the project. A.J.G., M.Z., R.S. and A.R.G. have written the manuscript. All authors have read and approved the final manuscript.

